# Magnetic Levitation and Sorting of Neoplastic Circulating Cell Hybrids

**DOI:** 10.1101/2022.11.03.515127

**Authors:** Kaitlyn Liang, Sena Yaman, Ranish K. Patel, Michael S. Parappilly, Brett S. Walker, Melissa H. Wong, Naside Gozde Durmus

## Abstract

Circulating hybrid cells (CHCs) are a novel, rare cell population that harbor tumor and immune cell phenotypes and genotypes and are detectible in peripheral blood. Several recent reports implicated CHCs in the metastatic cascade and found their enumeration to provide better prognostic value than conventionally-defined circulating tumor cells (CTCs). However, methods for isolation and enrichment of CHCs are not well-studied or established. Here, we developed an ultrasensitive, antigen-independent platform leveraging the principles of magnetic levitation for the detection and isolation of disseminated neoplastic CHCs. For the first time, we demonstrate that CHCs can be magnetically focused to different levitation heights, under various paramagnetic conditions using a static levitation system, and we quantified the biophysical properties of CHCs (i.e., levitation heights). In addition, we investigated whether magnetic levitation approach can be combined with the affinity-based strategies to enrich CHCs under the magnetic field. Using clinical samples from breast and colorectal cancer patients, we demonstrated that neoplastic cells can be sorted with a magnetic levitation-based sorting device, without relying on any surface markers. Overall, we demonstrated the feasibility of the magnetic levitation method for unbiased enrichment of rare neoplastic-immune hybrid cells from peripheral blood specimens from cancer patients. This approach can be expanded to more clinical samples and cancer types to unprecedentedly explore the biology of rare neoplastic cells and develop metastasis-tailored therapies broadly impacting personalized and precision clinical treatments.

## INTRODUCTION

A greater understanding of disseminated neoplastic cell biology is required for advances in the early detection of cancer and the identification of metastatic risk. One avenue that holds great promise is in developing platforms to isolate predictive biomarkers, such that these analytes can provide a non-invasive information for tumor burden. Further, isolation of non-invasive analytes, such as circulating neoplastic cells, will also provide a greater biologic understanding of their contribution to disease progression^1–6^. For instance, circulating tumor cells (CTCs) that are disseminated from primary tumors into the peripheral blood of cancer patients^13–17^ correlate with metastatic disease in most tumor types. However, their rarity complicates their value as reliable biomarkers in various cancer types and limits their value in providing biologic insight into mechanisms underlying tumor progression (i.e., 3 CTC/7.5 mL of blood in metastatic pancreatic cancer patients)^7^. Recently, a novel neoplastic hybrid cell population that harbors genotypes and phenotypes of two different cell lineages (i.e., immune cells and neoplastic cells; called circulating hybrid cells, CHCs) was identified^8–11^. These newly identified neoplastic hybrid cell populations, CHCs, outnumber the conventionally-defined CTCs in late stage disease (i.e., 10 times greater CHCs than CTCs in metastatic patients)^8^. CHCs have better prognostic value than CTCs, as their numbers correlated with stage and survival in pancreatic cancer, whereas CTC numbers did not^2^. Interestingly, CHCs also represent heterogeneous subpopulations, some of which may have high potential as effectors of metastatic tumor seeding. Thus, the detection and characterization of heterogeneous phenotypes of disseminated neoplastic CHCs in blood is critical for unraveling non-invasive clinical markers in cancer. Isolation of non-conventional neoplastic cells from peripheral blood will facilitate the development of new liquid biopsy tools for the detection, prognosis, and monitoring of disease progression; ultimately impacting patient outcomes.

Most of the available neoplastic cell isolation technologies depend on expression of known surface proteins and suffer from low throughput. The presence of heterogeneous subtypes of disseminated neoplastic cells (i.e., CTCs with EMT-like phenotypes and CHCs with enhanced metastatic potential) requires an unbiased method for separating these cells from blood without relying on specific antibodies, such as epithelial cellular adhesion molecule, EpCAM. There are various promising technologies utilizing the negative selection of white blood cells (WBCs) with magnetic beads to enrich CTCs. However, these methods are limited due to the high number of contaminating non-specific WBCs (order of 10^4^), confounding downstream analysis, and the exclusion of CHCs that harbor CD45 expression. In addition, while CHCs are more abundant in peripheral blood, their physical and phenotypic similarities to WBCs make their isolation and analysis a challenge^20^ (**Supplementary Figure 1**). Further, these devices operate at low flow rates, limiting sample processing. Our studies showed that the average CHC yield of a size-based sorting method was 4.03% ±4.17 (**Supplementary Figure 2**). Thus, there is a need for better technology platforms that isolate and enrich CHCs for enumeration and identification.

Here, we developed and optimized the efficacy of an ultrasensitive, antigen-independent platform leveraging the principles of magnetic levitation for the detection and isolation of disseminated neoplastic CHC populations (**Figure 1**). First, we showed that CHCs can be levitated and magnetically focused to different levitation heights; under various paramagnetic conditions using a static levitation system. We quantified the biophysical properties of CHCs (i.e., density and levitation heights). In addition, we investigated whether magnetic levitation approach can be combined with the affinity-based strategies (i.e., CD45 nanobeads) to isolate and enrich CHCs under the magnetic field. Using clinical specimen from cancer patients, we demonstrated that neoplastic cells can be sorted with a magnetic levitation-based sorting device; without relying on any surface markers. Overall, we demonstrated the feasibility of the magnetic levitation method for sorting rare CHCs from whole blood samples. When fully developed and validated with additional clinical samples and across various cancer sites, this approach has potential for unprecedented options to probe the biology of disseminated neoplastic cells, which may lead to metastasis-tailored therapies that broadly impact personalized and precision oncologic treatment paradigms. Further, application of an unbiased, disseminated tumor cell isolation platform may impact the evaluation of disease burden in cancer patients with potential to be deployed in the early detection setting, or for evaluation of treatment response.

**Figure 1.**
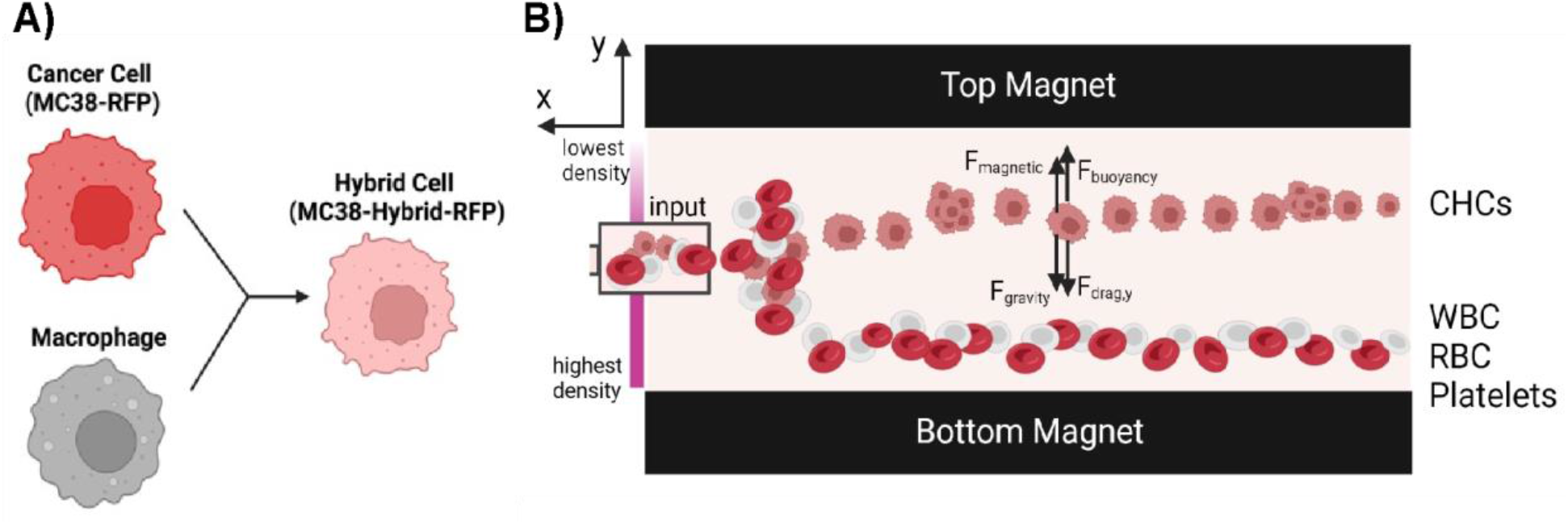
Magnetic levitation of circulating hybrid cells. **A)** Colorectal cancer cells (MC38 cells, RFP-labeled) and macrophages (Mϕ) spontaneously hybridize *in vitro* and form colorectal cancer hybrid cells (MC38 hybrid cells, RFP-labelled). **B)** Circulating cell fusion hybrids (CHCs) can be levitated under a magnetic field. In this magnetic configuration, magnetic forces acting on the CHCs counteract gravity. CHCs can be detected from the healthy blood cells (i.e., white blood, red blood cells, platelets) based on their different levitation profiles. F_magnetic_, F_buoyancy_, F_gravity_ and F_drag_ represent magnetic, buoyancy, gravity, and drag forces acting on the cells, respectively.

## RESULTS AND DISCUSSION

Cell fusion or the functional acquisition of features from a second lineage (i.e., generation of a hybrid cell) benefits the evolving cancer cell population, as the hybrid cells inherit genotypic and phenotypic characteristics of both two cell lineages to survive under selective pressure. Several reports present evidence that macrophages are an important partner to neoplastic cells in this process resulting in hybrid cells that retain tumor-initiating capacity and gain macrophage migratory capabilities^12–20^. Cell hybrids are identified in cell culture, detectible in murine cancer models, and found in human pancreatic ductal adenocarcinoma (PDAC) patients in their primary tumor, in peripheral blood as CHCs, and in metastatic lesions.^8^ To understand the magnetic levitation profiles of malignant cell hybrids, we evaluated a published and characterized *in vitro*-derived hybrid cell line^8^. MC38 hybrid cells were generated through spontaneous fusion between murine colorectal cancer cells and murine primary bone marrow-derived macrophages (**Figure 1a**). MC38 hybrid cells were levitated at 30 mM, 50 mM, and 80 mM paramagnetic medium for 20 minutes (**Figure 2**). Dotted black lines indicate approximate ranges of levitation heights of MC38 hybrid cells under each paramagnetic condition (**Figure 2a-c**). Hybrid cell levitation heights correlated with changes in the paramagnetic medium concentration, consistent with previous experiments and literature^21–25^. For instance, the levitation height of MC38 hybrid cells was 320 ± 29 μm at 30 mM paramagnetic media (**Figure 2a**), and their heights increased to 422 ± 19 μm (**Figure 2b**) and 468 ± 27 μm (**Figure 2c**), at 50 mM and 80 mM paramagnetic media, respectively.

**Figure 2.**
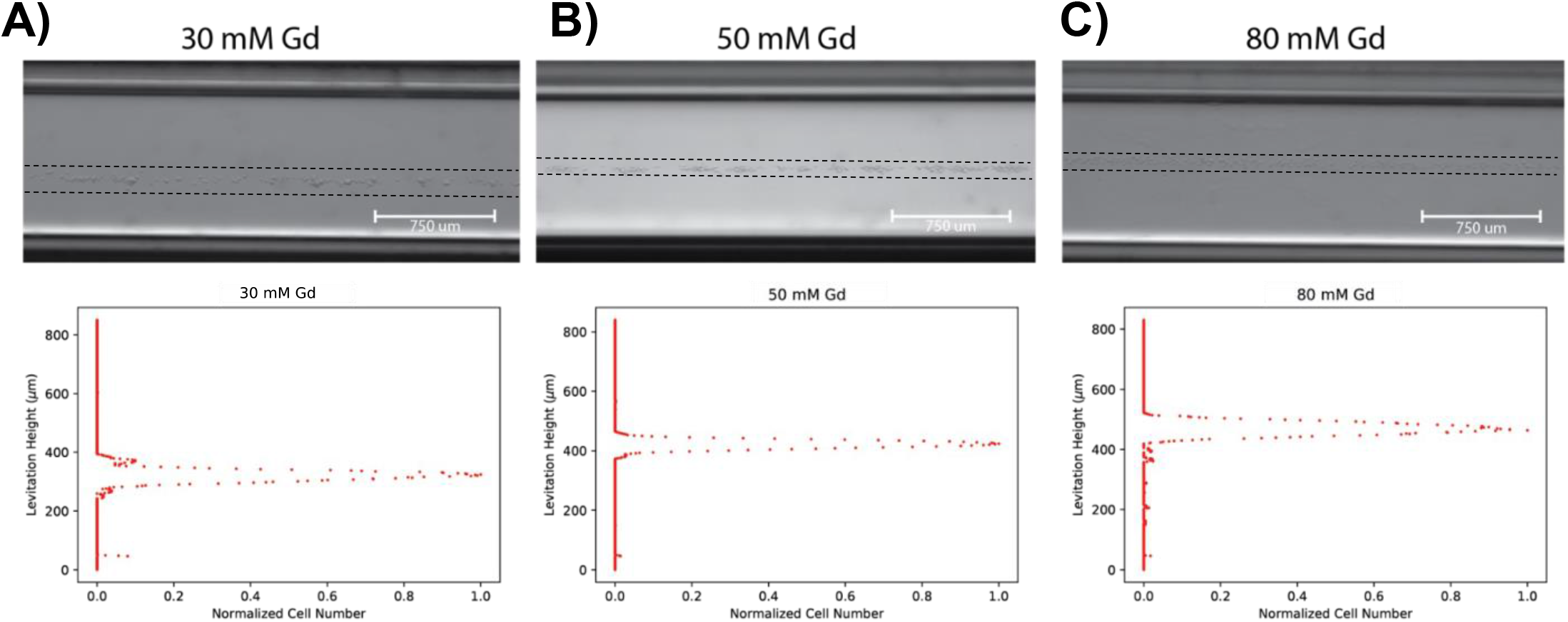
Characterization of magnetic levitation profiles of a hybrid cell line at different paramagnetic conditions. Static levitation bands of mouse MC38 hybrid cells levitated at various paramagnetic medium concentrations, such as **A)** 30 mM, **B)**50 mM, and **C)**100 mM. As molarity increases, levitation height increases, as cells are magnetically focused into the less dense regions of the capillary. Dashed lines signify levitation ranges of cell populations, indicating that diminishing heterogeneity is associated with increased paramagnetic medium concentration.

Next, we levitated and quantified the levitation profiles of mouse peripheral blood mononuclear cells (mPBMCs) at different paramagnetic conditions. In these experiments, we have worked with frozen stocks of mPBMCs. mPBMCs cells were levitated at 30 mM, 50 mM, and 80 mM paramagnetic medium for 20 minutes (**Figure 3**). Compared to the murine hybrid cell line, mPBMCs had a lower levitation height at all the paramagnetic conditions tested (**Figure 3a-c**). mPBMC levitation heights also correlated with changes in the paramagnetic medium concentration, consistent with previous experiments and literature^21–25^. For instance, the average levitation height of mPBMCs cells was 226 ± 58 μm at 30 mM paramagnetic media (**Figure 3a**), and their heights increased to 295 ± 75 μm (**Figure 3b**) and 400 ± 84 μm (**Figure 3c**), at 50 mM and 80 mM paramagnetic media, respectively. According to the literature, dead cells or cell debris have a higher density compared to the live cells, thus they levitate at lower positions in the magnetic levitation system^21,23^. For all the samples tested, the wider distribution of mPBMCs relative to the freshly CHC population could be a result of dead cells or cell debris generated from the freeze thawing process during sample preparation.

**Figure 3.**
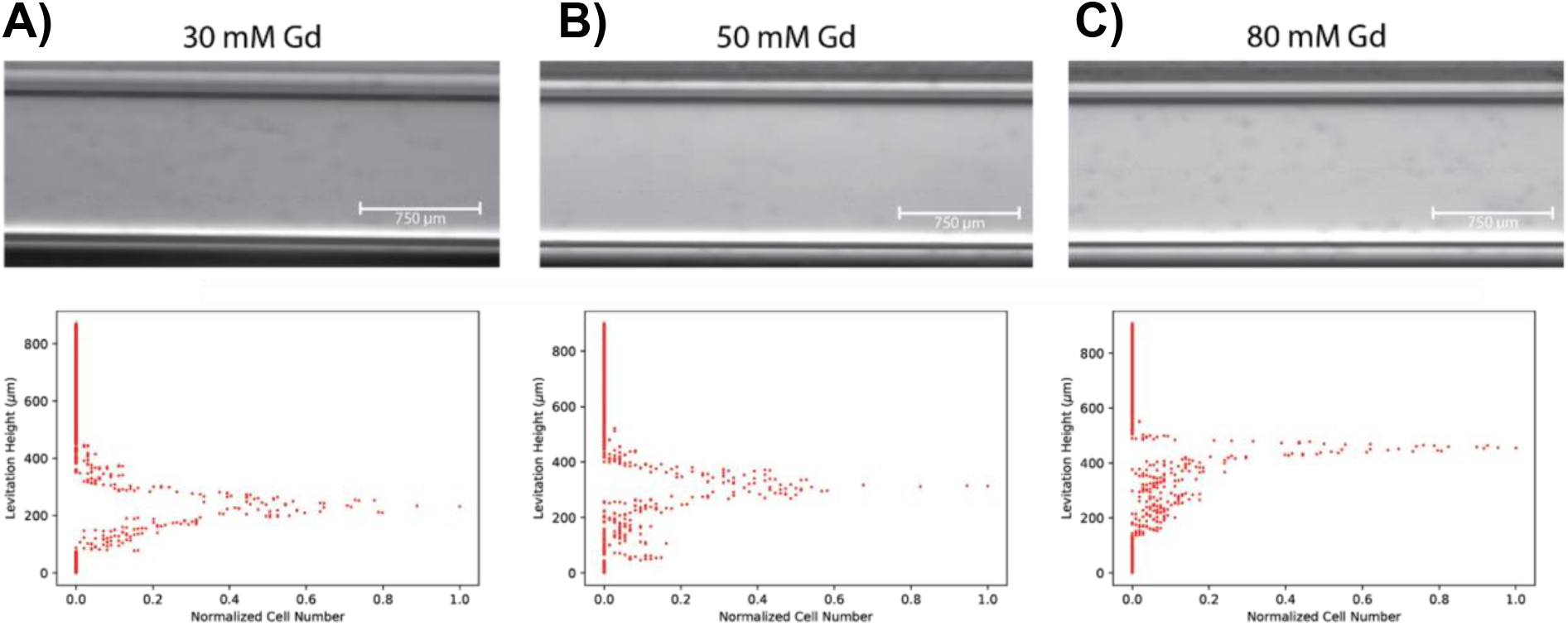
Characterization of magnetic levitation profiles of mouse peripheral blood mononuclear cells (mPBMCs) at different paramagnetic conditions. Static levitation bands of mPBMCs cells levitated at various paramagnetic medium concentrations: **A)** 30 mM, **B)** 50 mM, and **C)** 80 mM).

We next investigated whether the magnetic levitation approach can be combined with the affinity-based strategies to enrich hybrid cells under the magnetic field. For instance, CD45-expression distinguishes the CHCs (and hybrid cells) from previously defined circulating tumor cells (CTCs) that lack any immune cell expression. On the other hand, CD45 expression levels within hybrid cells (or patient isolated CHCs) is lower compared to the immune cells (i.e., PBMCs). Levitation profile of biological moieties (i.e., cells or proteins) can be manipulated upon the binding of magnetic nanobeads to the surface, thus by fine-tuning its overall magnetic susceptibility^26,27^. We investigated whether magnetic levitation could differentiate hybrid cells and PBMCs based on the minute differences in their surface marker expression (i.e., CD45) by manipulating their magnetic susceptibility in the presence of magnetic beads and focusing them into different levitation positions (**Figure 4**). We levitated the MC38 hybrid cells and mPBMCs in the presence of different CD45 microbeads concentrations (i.e., 1 μl, 5 μl and 10 μl microbeads). As the microbeads are magnetic, we hypothesized that upon their binding to the surface of mPBMCs, they will cause an instantaneous shift in their levitation profile as they have a higher CD45 expression, compared to the MC38 hybrid cells (**Figure 4a**). In the absence of CD45 microbeads, we observed that MC38 hybrid cells and mPBMCs levitated at similar positions after 10 minutes, as they did not have enough time to come to their own unique equilibrium positions which takes about 20 minutes (**Figure 4b**, control). After the addition CD45 microbeads, we showed that mPBMC levitation band started to disappear as most of mPBMCs (higher CD45 expression) were attached to the magnets upon binding of CD45 microbeads to their surface (**Figure 4b**, 1 μl CD45 microbeads, mouse). On the other hand, MC38 hybrid cells (lower CD45 expression) continued to levitate, and they formed a broader levitation band reflecting the heterogeneity of surface expression levels of each individual hybrid cell. As the concentration of microbeads increased, both MC38 hybrid cells (lower CD45 expression) and mPBMCs could no longer be levitated due to strong magnetic attraction; overcoming the paramagnetism of the surrounding medium and attached to the magnets (**Figure 4b**, 5 μl CD45 microbeads, mouse and 10 μl CD45 microbeads, mouse). Thus, we demonstrated that concentration of magnetic beads can be fine-tuned to detect very minute differences in surface marker expression levels between CHCs and immune cells. This approach could be further applied to enrich CHCs and deplete PBMCs to increase the purity of sorted samples for downstream genomic and transcriptomic characterization.

**Figure 4.**
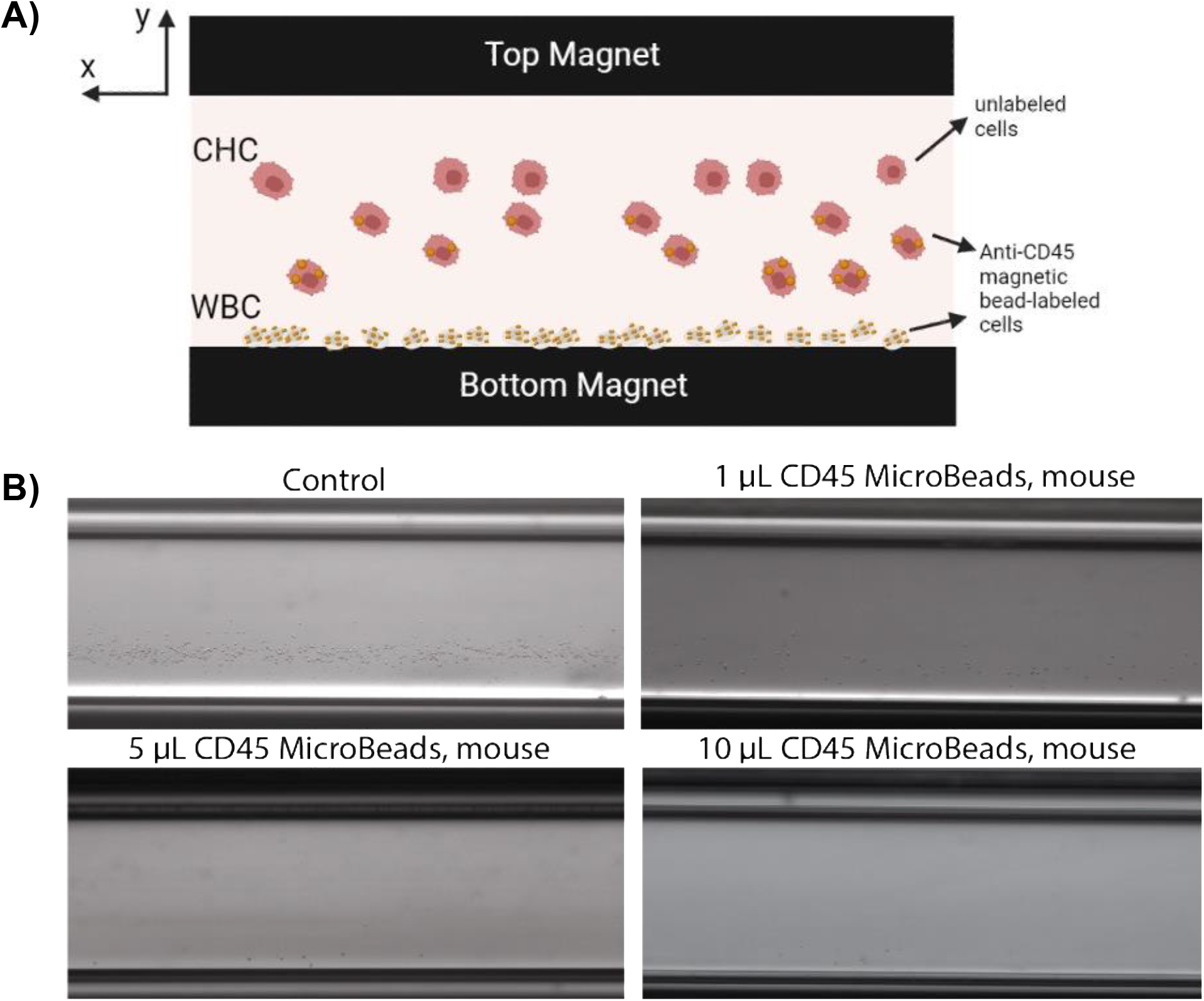
Optimization of white blood cell depletion and hybrid cell line enrichment with CD45 magnetic beads. **A)** Schematic of magnetic levitation of CHCs in the presence of magnetic CD45 beads. **B)** Optimization of PBMC depletion and hybrid cell line enrichment with different concentrations of CD45 magnetic beads in the static magnetic levitation system.

Next, we performed a feasibility study to investigate if CHCs can be sorted from clinical peripheral blood specimens, independent of their size or surface marker expression, using magnetic levitation (**Figure 5**). An early-stage breast cancer specimen and late-stage colorectal cancer specimen were processed with a flow-based magnetic levitation system. Briefly, PBMCs were isolated from whole blood using Ficoll-Paque as previously described^28^. PBMCs were diluted such that a 1 mL sample was sorted within the flow-based magnetic levitation system in a 30 mM paramagnetic medium. Cells that levitated above the PBMC levitation band were collected at the top collection port and immunostained for CHC-specific proteins to validate their identity. The PBMC levitation band was also collected from the bottom collection port, to determine if CHCs with greater immune identity might segregate with the PBMCs (**Figure 5a**). CHCs were identified as co-expressing the pan-leukocyte protein antigen, CD45, and either cytokeratin or EPCAM. We demonstrated that label-free, flow-based magnetic levitation system can detect and sort CHCs to the top port, for both breast cancer and colorectal cancer specimens. The bottom port was also stained and analyzed for CHCs. Interestingly, we detected presence of CHC subtypes with greater immune identity and phenotype more similar to an immune cell (resulting in a higher density and lower levitation height) at the bottom port. Thus, flow-based magnetic system can be potentially used to investigate the heterogeneity of CHCs in patient samples.

**Figure 5.**
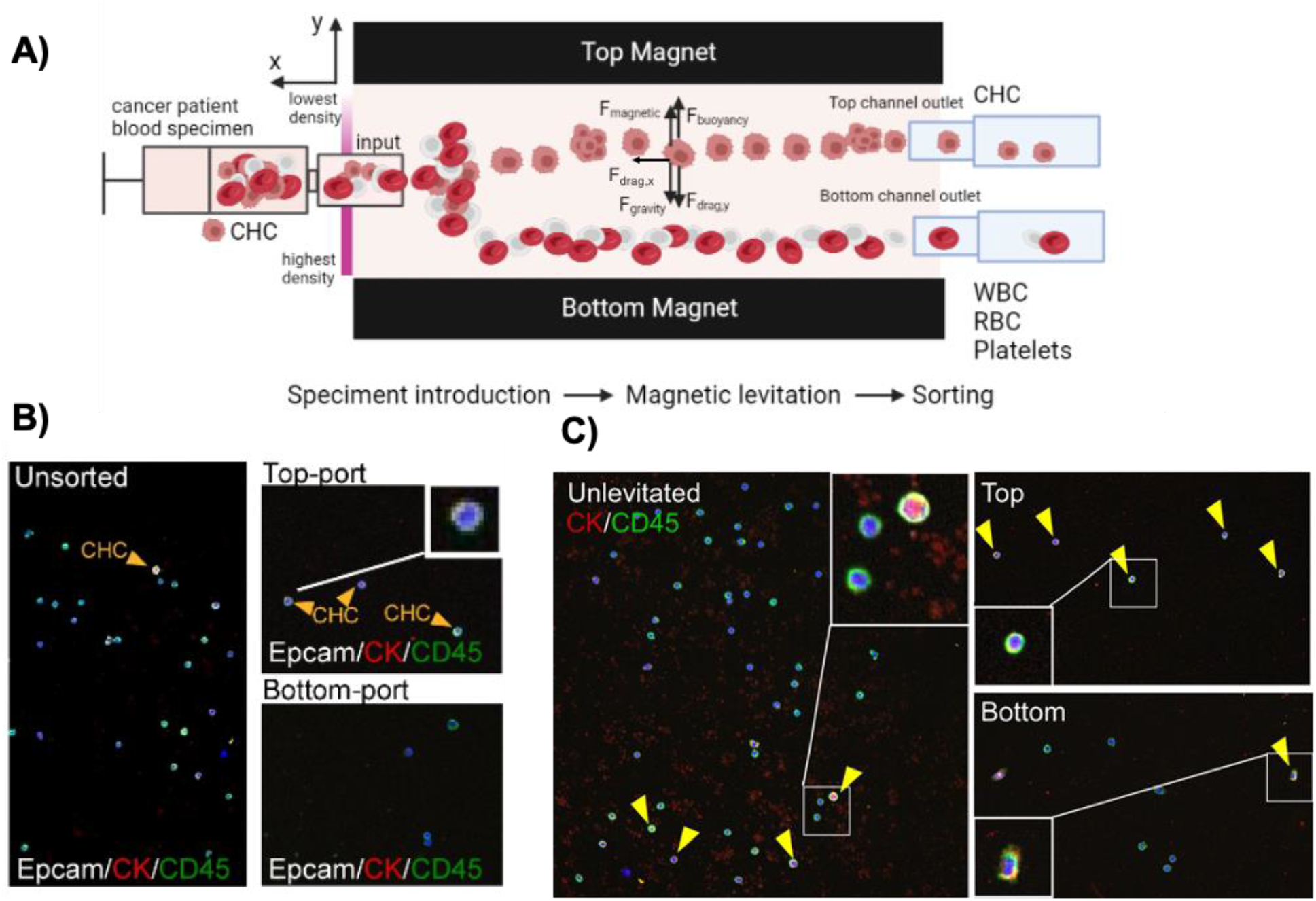
CHC enumeration and isolation from patient blood samples using a flow-based magnetic levitation system. **A)** Mechanism behind the label-free isolation technology. RBCs and PBMCs (higher density) are collected at bottom outlet, whereas CHCs (lower density) are collected at top outlet. Fmagnetic, Fbuoyancy, Fgravity and Fdrag represent magnetic, buoyancy, gravity, and drag forces acting on the cells, respectively. **B)** CHCs (yellow arrowhead) are isolated using levitational sorting, shown in “top-port” image. **C)** CHCs (yellow arrowhead) with more immune identity can be collected at the “bottom port”.

In summary, our study, for the first time, demonstrated that CHCs, a novel type of circulating neoplastic cells, can be levitated under a magnetic field. We demonstrated the feasibility of the magnetic levitation method for unbiased enrichment of rare cell fusion hybrids from cancer patient’s blood specimen. When validated with more clinical samples and cancer types, this label-free, magnetic-levitation approach represents an unprecedented option for understanding the biology of rare neoplastic cells and developing metastasis-tailored therapies that broadly impact personalized and precision oncologic treatment strategies. Further, application of an unbiased, disseminated tumor cell isolation platform may impact the evaluation of disease burden in cancer patients with potential to be deployed in the early detection setting, or for evaluation of treatment response.

## MATERIALS AND METHODS

### Levitation Device Design and Operation

The magnetic levitation device consists of laser-cut pieces of polymethyl methacrylate (PMMA), two permanent N52 neodymium bar magnets (K&J Magnetics) with the same poles facing each other, and custom-made mirrors^21–24^. A channel (1 mm in height) was placed between the magnets for static measurements. Angled side mirrors were used for real-time imaging and to fine-tune the magnetic focusing of cells. To sort the target cells, a flow-based magnetic levitation system was used. The flow-based system consisted of an inlet and two outlet ports (i.e., the top port collecting cells of interest less dense than the healthy blood cells, and the bottom port collecting the white blood cells). The samples can be introduced into the system by syringe pumps. In all flow experiments, a 2:1 flow ratio was maintained between the top and bottom ports.

### Cell Culture for mouse cancer cells and mouse hybrid cells

Murine MC38-RFP adenocarcinoma cells and MC38 hybrid cells (MC38-macrophage hybrid cells)^8^ were cultured in Dulbecco’s modified eagle medium (DMEM) (Thermo Fisher) supplemented with 10% FBS, 2mM glutamine, 0.1 mM nonessential amino acids, 1mM sodium pyruvate, 10 mM HEPES, 50 μg/mL gentamycin sulfate, 1% penicillin/streptomycin (Invitrogen Corp.). Cells were grown at 37 °C and 5% CO_2_in a humidified atmosphere.

### Magnetic levitation of mouse hybrid cells and mouse PBMCs

MC38xY01 cells and mouse PBMCs were diluted to 50,000 cells/mL in a 100 μL sample. Paramagnetic levitation solutions were prepared with non-cytotoxic, non-ionic, chelated gadolinium (Gd) ions, in DMEM cell culture media. Gadavist is an FDA-approved, human-injectable, non-toxic magnetic resonance imaging (MRI) contrast reagent. In some cases, calcein staining was used to distinguish MC38xY01 cells. PBS is used as the background medium. The Miltenyi Biotec CD45 MicroBeads (mouse) were attached by incubating the beads with the cells for at least 30 minutes at 4 °C before the addition of Gadavist. For static levitations, 30 μL of the sample was loaded into the capillary, and the capillary is inserted into the static levitation device. A 10-minute period is needed for the cells to equilibrate at its levitation height. Levitation height profiles were imaged and analyzed with inhouse developed image analysis software written on MATLAB, as described below.

### Magnetic levitation Height Analysis

A MATLAB program for automatic digital image processing was developed. The image datasets were acquired using the ZenPro2 software (Zeiss) and were imported into the code in “.tiff” format. The pixel heights corresponding to the top and bottom of the channel are measured. A sobel edge filter is applied to the image by convolving two 3×3 kernels representing the approximate derivatives in the horizontal and vertical directions. The result is the outlined shadows of the cell. Vertical and horizontal line additions and dilation will create a circular shape representing the cells. Closed objects are filled and eroded to create a more accurate signal of the cells. Tilting is corrected, and the number of bright pixels at each pixel height in the channel is counted. Once normalized, the levitation height is determined.

### Sorting of CHCs from mouse blood using commercial Parsortix system

The Parsortix PC1 Clinical System is a commercially used device for the capture and harvest of circulating tumor cells (CTCs) from patient blood samples through the use of microfluidics. CTCs and CHCs are morphologically similar, so this device was used to set a commercial standard for CHC sorting. A 200 μL sample containing 1:10 dilution mouse PBMCs (stock solution is 1 million cells/mL), and 1,000 cells/mL concentration of calcein-stained MC38xY01 is prepared. The sample is then run through the Parsortix system where the MC38xY01 cells will theoretically get caught in the filtration cassette due to their larger size and lower compressibility. The cells were then eluted and examined under a microscope to determine the sorting specificity and sensitivity.

### Levitational sorting of CHCs from clinical blood samples

Deidentified patient peripheral blood samples (one an early-stage breast cancer sample and one late-stage colorectal cancer sample) were retrieved from Oregon Health & Science University (OHSU). Informed consent was obtained from all subjects. All experimental protocols were approved by the OHSU Institutional Review Board. Whole blood samples were processed immediately after blood collection. PBMCs were separated using Ficoll-Paque, as previously described^28^. PBMCs were diluted in PBS at a 1:10 ratio, and then were levitated and sorted in a 30 mM paramagnetic medium. Cells that levitated above the PBMC levitation band were collected at the top and bottom collection ports then immunostained with CHC-specific antibodies (against EpCAM and CD45) for validation.

### Immunohistochemistry for evaluation of enriched CHCs

After levitational sorting, collected cells were first treated with a 5% bovine serum albumin solution, followed by TrueBlack Lipofuscin Autofluorescence Quencher (Biotium), and Image-iT FX Signal Enhancer (Invitrogen), then stained with fluorescent-conjugated antibodies for pan-cytokeratin (CK; eBioscience, clone: AE1/AE), and CD45 (Biolegend, clone:HI30), and counterstained with the nuclear dye, DAPI. Each sample was processed with unstained and isotype controls (eBioscience). Specimens were digitally imaged with a Zeiss AxioScanner.

## Acknowledgements

N.G.D. acknowledges support from the Career Award at the Scientific Interface (CASI) from the Burroughs Wellcome Foundation (BWF). N.G.D. acknowledges support from the McCormick and Gabilan Faculty Award from Stanford University. M.H.W. acknowledges support from Department of Defense (W81XWH-18-1-0621) and the National Institutes of Health/National Cancer Institute (CA260196-0A1). M.S.P is supported by the National Institutes of Health (TL1TR002371).

## Supplementary Figures

**Supplementary Figure 1.**
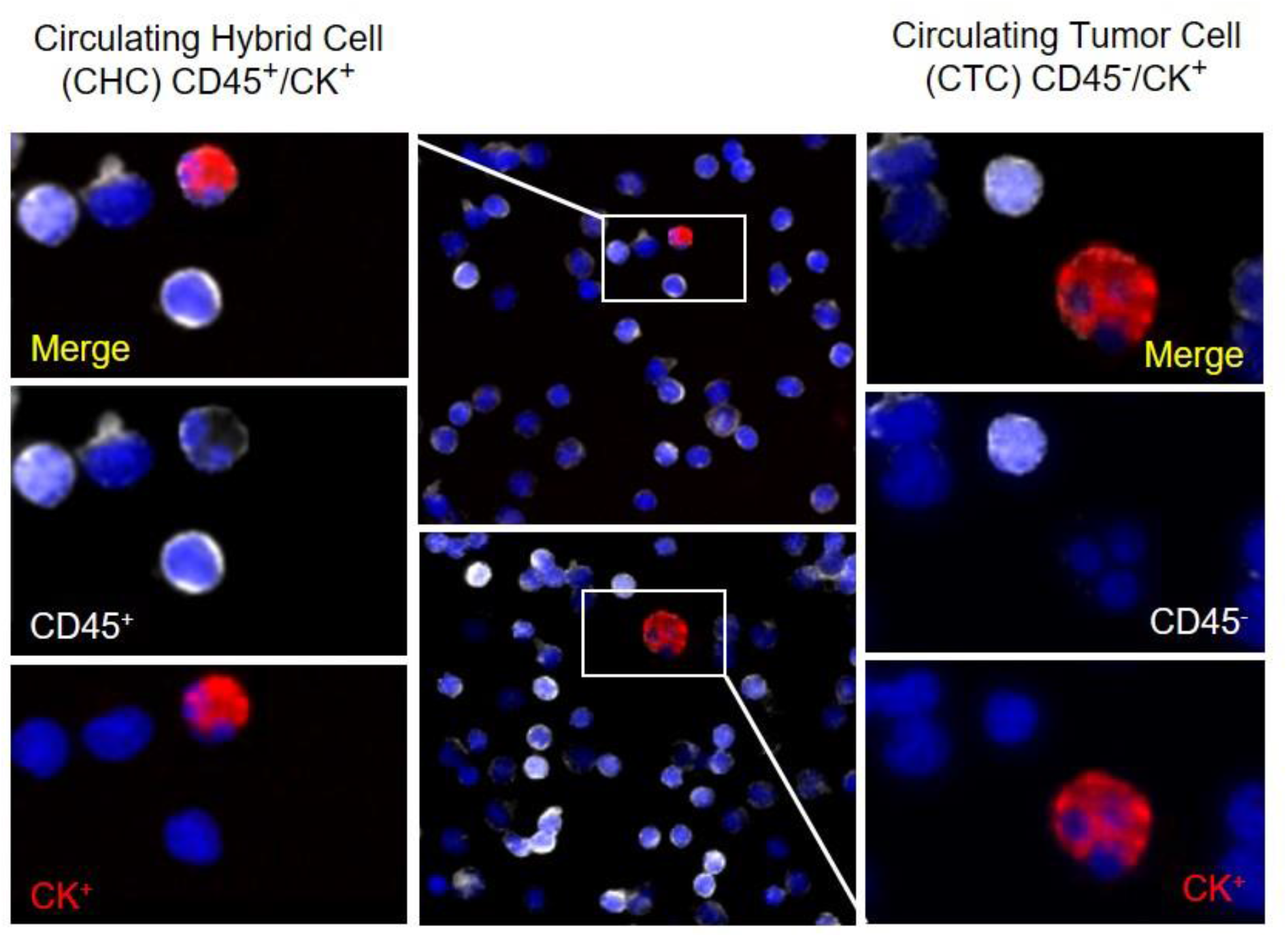
Circulating hybrid cells (CHC) and circulating tumor cells (CTC) are detected in the peripheral blood of patients with colon adenocarcinoma. CHCs are identified by co-expression of CD45 (white) and cytokeratin (CK; red). CTCs lack CD45 expression while expressing CK and are larger than CHCs. Nuclear DAPI staining is blue.

**Supplementary Figure 2.**
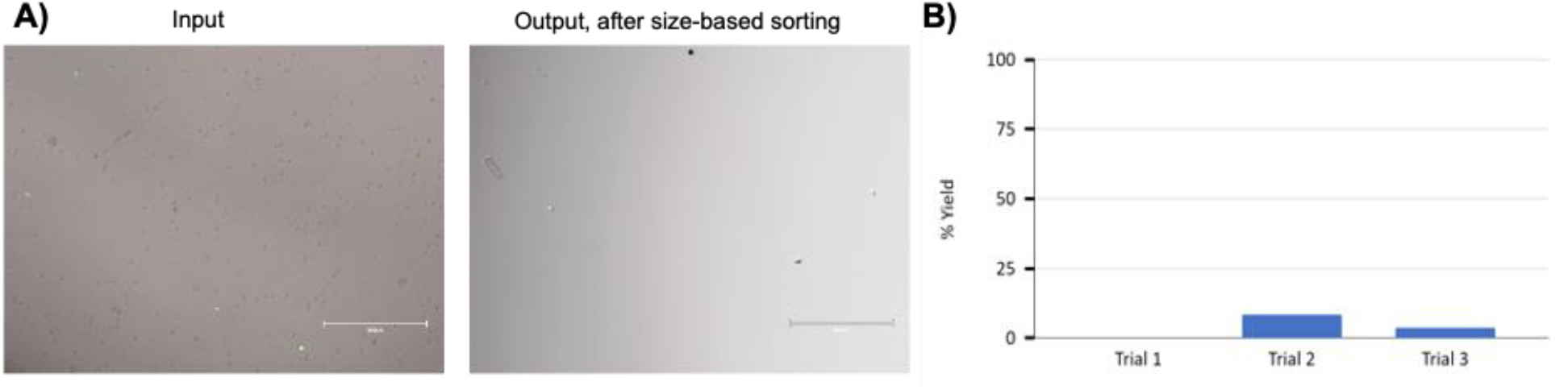
Sorting of MC38xY01 hybrid cells from mouse PBMC using a size-based sorting strategy. **A)** Representative pictures of the input sample (calcein-stained hybrid cells+ mPBMCs) and output sample after size-based sorting. The small dark circles are the mPBMCs, and any irregularly shaped object is most likely cell debris. **B)** The sorting yield after each trial. The average yield of a size-based sorting method was %4.03±4.17.

## REFERENCES

1 Aceto, N. et al. Circulating tumor cell clusters are oligoclonal precursors of breast cancer metastasis. Cell 158, 1110–1122 (2014).

2 Hou, J.-M. et al. Circulating tumor cells as a window on metastasis biology in lung cancer. The American journal of pathology 178, 989–996 (2011).

3 Yu, M. et al. Circulating breast tumor cells exhibit dynamic changes in epithelial and mesenchymal composition. science 339, 580–584 (2013).

4 Cho, E. H. et al. Characterization of circulating tumor cell aggregates identified in patients with epithelial tumors. Physical biology 9, 016001 (2012).

5 Molnar, B., Ladanyi, A., Tanko, L., Sréter, L. & Tulassay, Z. Circulating tumor cell clusters in the peripheral blood of colorectal cancer patients. Clin Cancer Res 7, 4080–4085 (2001).

6 DiPardo, B. J., Winograd, P., Court, C. M. & Tomlinson, J. S. Pancreatic cancer circulating tumor cells: applications for personalized oncology. Expert Rev Mol Diagn 18, 809–820, doi:10.1080/14737159.2018.1511429 (2018).

7 Gao, Y. et al. Clinical significance of pancreatic circulating tumor cells using combined negative enrichment and immunostaining-fluorescence in situ hybridization. J Exp Clin Cancer Res 35, 66, doi:10.1186/s13046-016-0340-0 (2016).

8 Gast, C. E. et al. Cell fusion potentiates tumor heterogeneity and reveals circulating hybrid cells that correlate with stage and survival. Sci Adv 4, eaat7828, doi:10.1126/sciadv.aat7828 (2018).

9 Powell, A. E. et al. Fusion between Intestinal epithelial cells and macrophages in a cancer context results in nuclear reprogramming. Cancer Res 71, 1497–1505, doi:10.1158/0008-5472.CAN-10-3223 (2011).

10 Rizvi, A. Z. et al. Bone marrow-derived cells fuse with normal and transformed intestinal stem cells. Proceedings of the National Academy of Sciences of the United States of America 103, 6321–6325, doi:10.1073/pnas.0508593103 (2006).

11 Dietz, M. S. et al. Relevance of circulating hybrid cells as a non-invasive biomarker for myriad solid tumors. Sci Rep 11, 13630, doi:10.1038/s41598-021-93053-7 (2021).

12 Adams, D. L. et al. Circulating giant macrophages as a potential biomarker of solid tumors. Proc Natl Acad Sci U S A 111, 3514–3519, doi:10.1073/pnas.1320198111 (2014).

13 Pawelek, J. M. & Chakraborty, A. K. Fusion of tumour cells with bone marrow-derived cells: a unifying explanation for metastasis. Nature Reviews Cancer 8, 377–386 (2008).

14 Clawson, G. A. et al. Macrophage-tumor cell fusions from peripheral blood of melanoma patients. PloS one 10, e0134320 (2015).

15 Clawson, G. A. et al. “ Stealth dissemination” of macrophage-tumor cell fusions cultured from blood of patients with pancreatic ductal adenocarcinoma. PLoS One 12, e0184451 (2017).

16 Manjunath, Y. et al. Circulating giant tumor-macrophage fusion cells are independent prognosticators in patients with NSCLC. Journal of Thoracic Oncology 15, 1460–1471 (2020).

17 Lizier, M. et al. Fusion between cancer cells and macrophages occurs in a murine model of spontaneous neu+ breast cancer without increasing its metastatic potential. Oncotarget 7, 60793 (2016).

18 Ding, J., Jin, W., Chen, C., Shao, Z. & Wu, J. Tumor associated macrophage× cancer cell hybrids may acquire cancer stem cell properties in breast cancer. PloS one 7, e41942 (2012).

19 Cao, M.-F. et al. Hybrids by tumor-associated macrophages× glioblastoma cells entail nuclear reprogramming and glioblastoma invasion. Cancer Letters 442, 445–452 (2019).

20 Zhang, L.-N., Huang, Y.-H. & Zhao, L. Fusion of macrophages promotes breast cancer cell proliferation, migration and invasion through activating epithelial-mesenchymal transition and Wnt/β-catenin signaling pathway. Archives of biochemistry and biophysics 676, 108137 (2019).

21 Durmus, N. G. et al. Magnetic levitation of single cells. Proc Natl Acad Sci U S A 112, E3661–3668, doi:10.1073/pnas.1509250112 (2015).

22 Puluca, N. et al. Levitating Cells to Sort the Fit and the Fat. Adv Biosyst, e1900300, doi:10.1002/adbi.201900300 (2020).

23 Chin, E. K., Grant, C. A., Ogut, M. G., Cai, B. & Durmus, N. G. CelLEVITAS: Label-free rapid sorting and enrichment of live cells via magnetic levitation. bioRxiv, 2020.2007.2027.223917, doi:10.1101/2020.07.27.223917 (2020).

24 Urey, D. Y., Chan, H. M. & Durmus, N. G. Levitational Cell Cytometry for Forensics. Adv Biol (Weinh) 5, e2000441, doi:10.1002/adbi.202000441 (2021).

25 Goreke, U., Bode, A., Yaman, S., Gurkan, U. A. & Durmus, N. G. Size and density measurements of single sickle red blood cells using microfluidic magnetic levitation. Lab Chip 22, 683–696, doi:10.1039/d1lc00686j (2022).

26 Yaman, S. & Tekin, H. C. Magnetic susceptibility-based protein detection using magnetic levitation. Analytical Chemistry 92, 12556–12563 (2020).

27 Shapiro, N. D., Soh, S., Mirica, K. A. & Whitesides, G. M. Magnetic Levitation as a Platform for Competitive Protein–Ligand Binding Assays. Analytical Chemistry 84, 6166–6172, doi:10.1021/ac301121z (2012).

28 Montoya Mira, J. et al. Label-free enrichment of rare unconventional circulating neoplastic cells using a microfluidic dielectrophoretic sorting device. Commun Biol 4, 1130, doi:10.1038/s42003-021-02651-8 (2021).

